# scDirect: key transcription factor identification for directing cell state transitions based on single-cell multi-omics data

**DOI:** 10.1101/2024.01.08.574757

**Authors:** Chen Li, Sijie Chen, Yixin Chen, Haiyang Bian, Minsheng Hao, Lei Wei, Xuegong Zhang

## Abstract

Cell state transitions are complicated processes that occur in various life activities. Understanding and artificially manipulating them have been longstanding challenges. Substantial experiments reveal that the transitions could be directed by several key transcription factors (TFs). Here we present scDirect, a computational framework to identify key TFs based on single-cell RNA-seq and ATAC-seq data. scDirect models the TF identification task as a linear inverse problem, and solve it with gene regulatory networks enhanced by a graph attention network. Through a benchmarking on a single-cell human embryonic stem cell atlas, we demonstrate the robustness and superiority of scDirect against alternative analysis methods on TF identification. We apply scDirect on various datasets, and scDirect exhibits high capability in identifying key TFs in cell differentiation and somatic cell conversion. Furthermore, scDirect can efficiently identify TF combinations for cell reprogramming, many of which have been experimentally validated. We envision that scDirect can utilize rapidly increasing single-cell datasets to identify key TFs for directing cell state transitions and may become an effective tool to facilitate cell engineering and regenerative medicine.

## 1 Introduction

Transitions between different cellular states are essential in various biological processes, such as cell differentiation, development, and disease. The widespread practice of cell engineering has successfully implemented transitions of different cellular states, including reprogramming of somatic cells to pluripotent stem cells, directional differentiation of pluripotent stem cells to somatic cells, and direct conversions between somatic cells^1-4^. These transitions are commonly implemented with forced expression of several specific transcription factors (TFs). It has been proved that a small number of key TFs is sufficient to direct cell state transitions^5-7^.

There are roughly 2,000 different TFs in humans^8,9^, composing a large space of plausible sets of TFs. It is time-consuming and labor-intensive to experimentally identify key TFs that can direct a certain cell state transition from the space. Researchers have developed a series of computational methods to help identify key TFs in cell state transitions^10-12^. Most of these methods are designed to predict TFs based on bulk omics data generated from a mixed population of cells or tissues. D’Alessio et al.^13^ proposed an entropy-based method to calculate a score for each TF from gene expression data. CellNet^14^ incorporates the gene regulatory network (GRN) structure inferred from gene expression and calculates a network influence score for each TF. The score represents how well the TF-perturbed GRN matches the target-cell-state GRN. As GRNs inferred from gene expression data may bring some false positive TF-target pairs^10^, methods like Mogrify^15^ introduce the external DNA-protein interactions from STRING^16^ to construct an experimentally validated GRN. Moreover, ANANSE^17^ builds GRNs from both gene expression and chromatin immunoprecipitation sequencing (ChIP-seq) data, and TFs are ranked according to their degree of regulating genes vital to cell reprogramming. Besides methods primarily based on gene expression, there are some other methods that only employed chromatin accessibility profiles achieving good performance^11,18,19^.

These bulk-data-based methods require abundant samples of cells with candidate states^14^, and are thus unusable for rare cell types or undefined cell states. The heterogeneity of the bulk data may also reduce the quality of TF identification^20^. Yet, the development of single-cell sequencing technologies^21-23^ has provided an unprecedent opportunity to facilitate the study of cell identity characterization, GRN inference of GRNs, and cell state transitions at the single-cell level. As for GRN construction, the methods based on both single-cell RNA sequencing (scRNA-seq) data and single-cell Assay for Transposase-Accessible Chromatin using sequencing (scATAC-seq) data exhibit better performance than previous methods based on single-omics data^24-26^. The constructed GRN can be utilized for TF perturbation, the inverse task of TF identification, which perturbs cells by the influence of given specific TFs’ variation propagating on the GRN^25^. Regarding TF identification in cell state transitions, some methods like differential expression analysis (DEA) and Dynamo can be employed as alternative solutions^27,28^, while these methods only output important genes during cell state transitions instead of key TFs responsible for directing cell state transitions.

To better leverage the significantly increasing single-cell datasets for TF identification, we present scDirect, a computational framework to identify key TFs in cell state transitions. scDirect models the TF identification task as a linear inverse problem, and leverage the graph attention network (GAT)-enhanced GRN inferred from single-cell multi-omics data to solve the alterations of total TFs. We benchmark scDirect on a TF-overexpression human embryonic stem cell (hESC) atlas^29^, and the results show a superior performance of scDirect. Then we apply scDirect on a mouse single-cell multi-omics dataset^23^, and known key TFs of different lineages of mouse hair follicle development are identified by scDirect. Furthermore, we curate 5 single-cell datasets of different experimentally validated reprogramming cases, and scDirect identifies most key TFs of these cases and outperforms existing methods. Moreover, scDirect could quantitatively identify feasible reprogramming TF combinations among thousands of possible combinations. Together, these results suggest scDirect is a reliable and effective tool to identify key TFs directing cell state transitions at the single-cell level.

## 2 Results

### 2.1 Overview of the scDirect framework

We developed scDirect as a computational framework for identifying key TFs directing cell state transitions from the source state to the target state (Fig. 1a). scDirect takes both scRNA-seq data and scATAC-seq data as inputs. The scRNA-seq data should contain cells of both the source and target states. The scATAC-seq data are not required to be simultaneously sequenced in the same cells of the scRNA-seq data but are best to be derived from similar tissues to guarantee the reliability. These data are first employed to construct a primary GRN with CellOracle^25^ which is proved to be a reliable tool for constructing linear GRNs (Methods). Here, scATAC-seq data provide the knowledge of regulatory directions from TFs to target genes, and scRNA-seq data are used to calculate the corresponding regulatory coefficients of TFs to target genes.

**Fig. 1.**
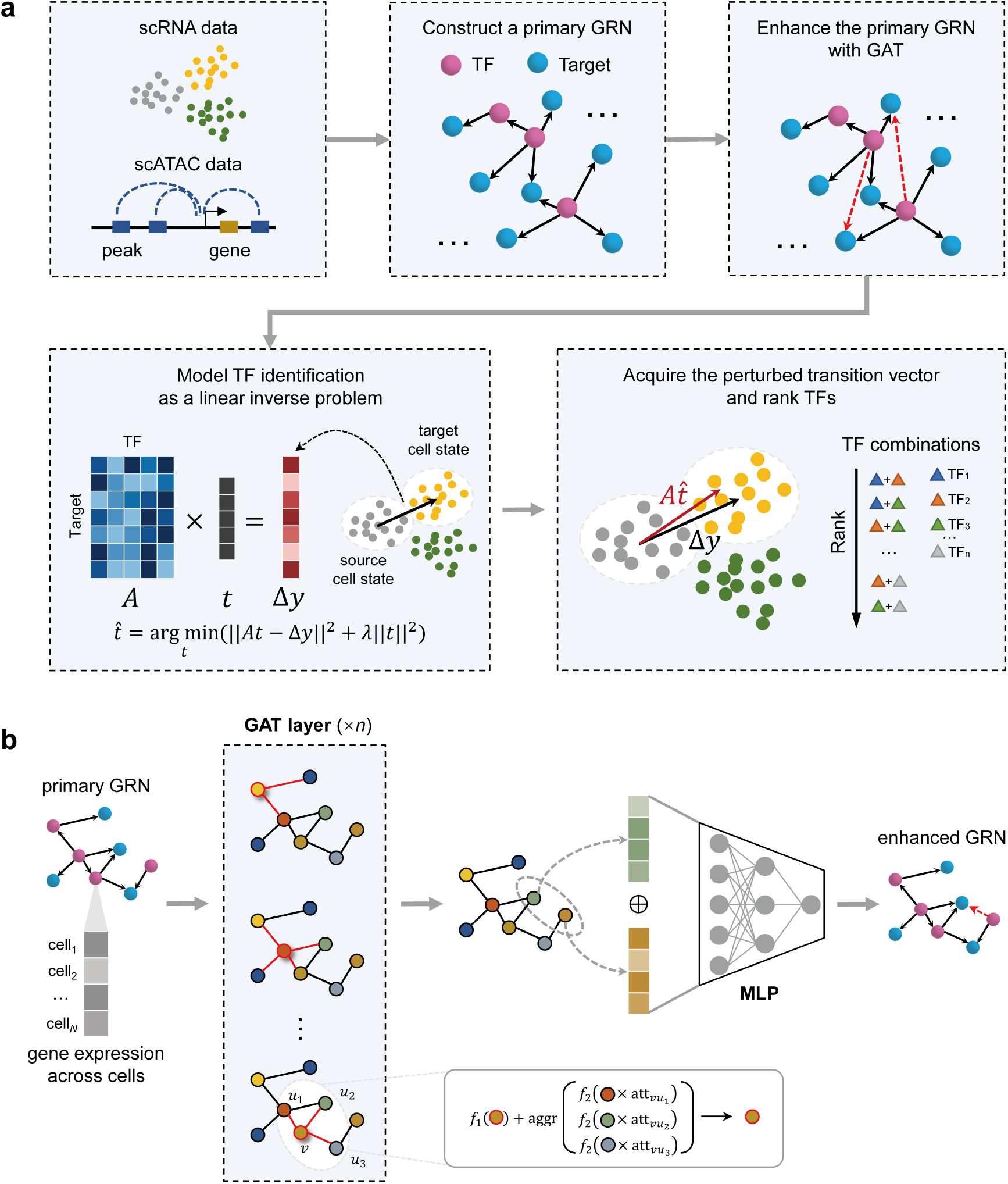
A graphical illustration of scDirect. **a** scDirect first constructs a primary GRN with scRNA data and scATAC data, and then it enhances the primary GRN with GAT. The TF identification task is modelled as a linear inverse problem and solved with Tikhonov regularization. Finally, scDirect uses the calculated expected alteration to get the directing score of each TF. **b** A graphical illustration of GAT recovering missing links. The normalized gene expression and the primary regulatory network comprise the input graph, and each node represents a gene. The whole model consists of the GAT encoder and the multilayer perceptron predictor. Multi-head attention mechanism is applied in the GAT layer to stabilize the learning process.

The dropout events of scATAC-seq data and the incomplete TF binding motif knowledge may lead to the missing of important TF-target links. Following the idea that graph neural networks can predict the interactions of genes^30,31^, scDirect employs a GAT model to recover missing TF-target links (Fig. 1b). Taking the primary GRN as the input training data, scDirect trains a GAT model and then applies the model to obtain additional putative TF-target pairs with high confidence (Methods). After GAT enhancement, scDirect acquires an enhanced GRN, which may have more complete regulatory relations and can better describe the whole transition process.

Subsequently, scDirect models the TF identification task as a linear inverse problem. With the GRN matrix ***A*** and the cell state transition vector *Δ****y***, scDirect solves the expected alteration vector 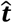 of TFs with Tikhonov regularization (Methods). Then, for each single TF or TF combination, the corresponding expected alterations are fixed to get a specific alteration vector 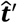. scDirect calculates the Pearson correlation coefficient (PCC) between 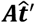 and Δ***y*** as the directing score, which assesses the importance of a single TF or a TF combination in directing the cell state transition. The final TF set is selected based on both expected alterations and directing scores with a same quantile threshold, and ranked by directing scores.

### 2.2 Graph attention networks enhance the gene regulatory signals

We adopted the GRN benchmark in CellOracle to validate the improvement of GAT model on the original GRN. Five tissues available both in the Tabula Muris scRNA-seq dataset^32^ and mouse sci-ATAC-seq atlas data^33^ were selected as the ground-truth datasets: heart, kidney, liver, lung and spleen. The ground-truth GRNs were curated using 1,298 chromatin immunoprecipitation followed by sequencing (ChIP–seq) datasets for totally 80 regulatory factors across these tissues^25^. The TF number of each ground-truth GRN ranges from 7 to 44, and the TF-target connection number ranges from 340 to 33,247. The scATAC-seq data was provided by mouse sci-ATAC-seq atlas^33^, and 13 samples across 5 tissues in the scRNA-seq dataset were utilized to conduct the GRN benchmark.

For each sample, we used CellOracle to calculate a primary GRN, and then GAT was applied to enhance the GRN. As for the GAT training, we performed a ten-fold cross-validation approach to improve the quality of the recovering TF-target links (Methods). As shown in Fig. 2a, the GAT prediction model gives the prediction result with a high area under the precision-recall curve (AUPRC) score on the test sets across all the 13 samples, which indicates the model robustly learns the regulatory relations between TFs and targets for each fold. For example, in sample Liver_2 and Spleen_0, the prediction model achieves AUPRC over 0.980 across all folds. The fold-average AUROC score ranges from 0.917 (Kidney_2) to 0.995 (Lung_0) across samples, which indicates GAT model is highly effective in capturing the regulatory signals of different tissues.

**Fig. 2.**
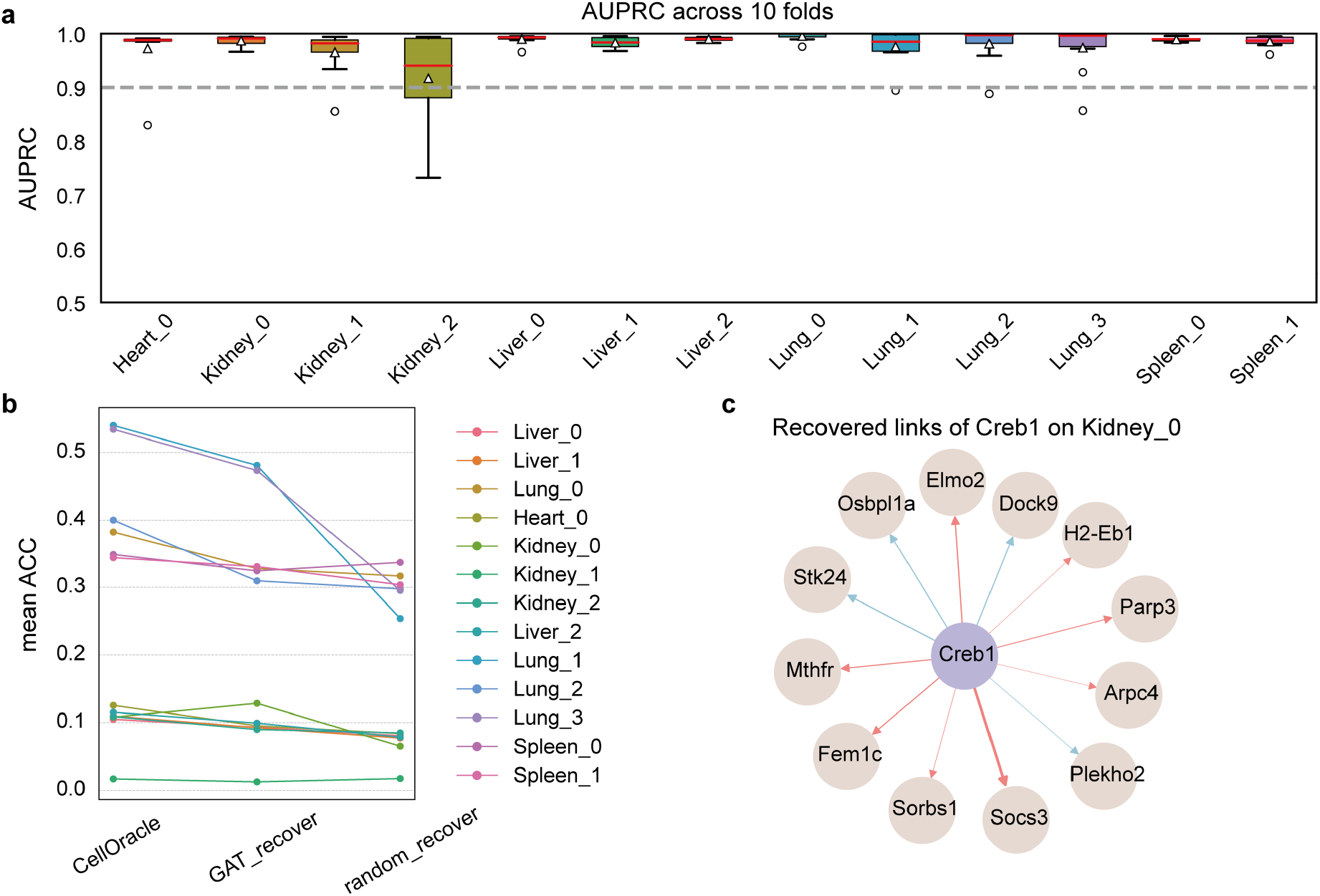
Graph attention network (GAT) module enhances the regulatory signal of original GRN. **a** Boxplots of area under the receiver operating characteristic (AUROC) on the test set across 13 samples. The box plots indicate the medians (centerlines), means (triangles), first and third quartiles (bounds of boxes). **b** Mean TF-target prediction accuracy comparison across CellOracle, GAT recovering, and random sampling. **c** GAT recovered network of Creb1. Ground-truth links are annotated in red.

To further investigate the GAT-recovered TF-target links, we calculated the accuracy (ACC) of predicted links compared to the ground-truth GRN for each TF in different samples. Then, we randomly sampled unconnected links, which number is equal to the number of predicted links. The sampled links could be regarded as the control group. The link prediction ACC comparison among CellOracle, GAT recovering and random sampling are illustrated in Fig. 2b. The performance of GAT-recovered links predominantly depends on the quality of original GRN. Among 11 of 13 samples, the predicted links exhibited higher mean ACC scores than the random links, which suggests that the GAT module recovers biologically meaningful regulatory links.

Moreover, we illustrated the recovered links of a specific TF Creb1, a kind of cyclic adenosine monophosphate (cAMP) responsive element modulator, in sample Kidney_0 (Fig. 2c). 8 of 12 links (red lines) are in the ground-truth kidney GRN, and Arpc4 and Mthrf are demonstrated to be targets of Creb1 in JASPAR predicted transcription factor targets dataset^34,35^. All the results demonstrate that the GAT module captures the regulatory relations and enhances the regulatory signals of the primary GRN.

### 2.3 Benchmarking TF identification on the TF-overexpression hESC atlas

To our knowledge, there is not a quantitatively benchmark of TF identification for single-cell datasets. We adopt a single-cell atlas built by Joung et al. to construct the benchmark of TF identification^29^. This atlas profiled hESCs infected with a lentivirus library to perform single TF overexpression in each cell. With TF overexpression directing hESCs to differentiate into different cell states, this atlas can be an experimentally validated ground truth to evaluate the key TF identification in cell state transitions. We selected cells of one source state and 15 target states to construct the benchmark dataset (Methods). Each target state contains a set of ground-truth key TFs, and the number of the ground-truth TFs ranges from 1 to 8 (Fig. 3a). For the benchmarking metric, we defined a TF identification score (TIS) to evaluate the identified TFs based on the ranks and amounts (Methods).

**Fig. 3.**
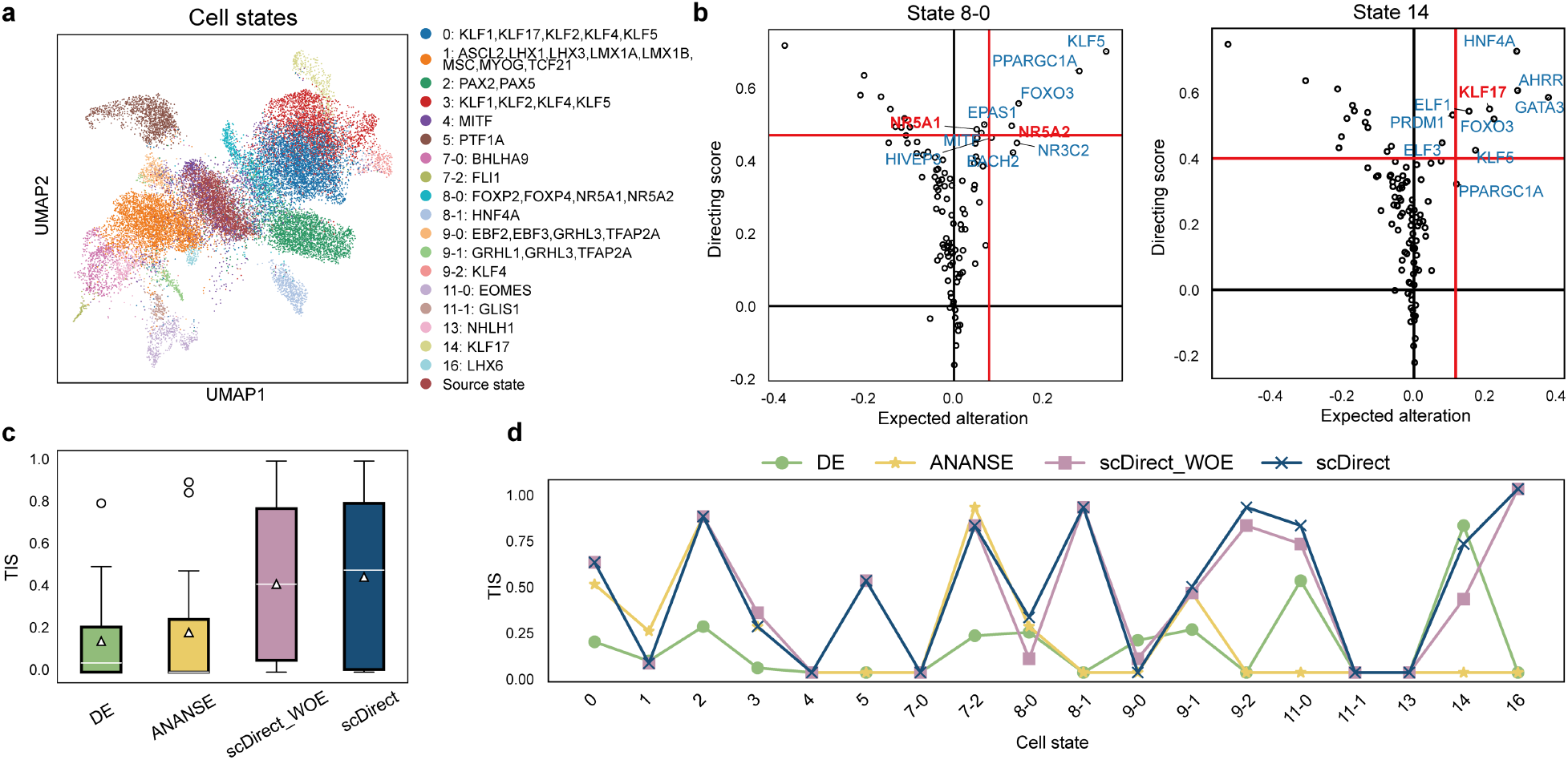
TF identification benchmarking on a single-cell human embryonic stem cell (hESC) atlas. **a** UMAP visualization of the single-cell hESC atlas. **b** scDirect TF identification plot on target state 8-0 and 14. Red lines are the quantile thresholds to filter 10 TFs. Key TFs are annotated in red. **c** TF identification score (TIS) comparison across 18 target states. The box plots indicate the medians (centerlines), means (triangles), first and third quartiles (bounds of boxes). **d** Line plots of TIS comparison across 18 target states.

We compared scDirect with some baseline methods. DEA was performed by the Wilcoxon rank-sum test. Existing methods for TF identification such as Mogrify and ANANSE are not practicable in the benchmarking as they require to collect datasets for specific cell states. Besides, the design for bulk data also hinders the application of these methods. To overcome the limitations, we replace the GRN inference part of ANANSE by CellOracle, and ANANSE is then used to calculate the importance score of each TF in the differential GRN of the target and source state. We also included scDirect without GAT enhancement (scDirect_WOE) to evaluate the effect of the GAT module. For each method, a list of ranked top 10 TFs is acquired to perform the benchmark.

As shown in Fig. 3c, scDirect significantly outperformed DEA and ANANSE and achieved higher TISs. scDirect achieved comparable or better performance than DEA in 16 of 18 target states except the target state 8-0 and 14 (Fig. 3d). In most target states, such as the target state 8-1 and 16, the ground-truth key TFs ranked high in scDirect identified TF list, while they did not appear in the DEA identified list. ANANSE achieved best performance in target state 1 and 7-2, while in most of remaining states it showed poor performance. It is reasonable of the instability of ANANSE because it ignores part information of TFs whose difference between target state and source state is not statistically significant. Compared to scDirect_WOE, scDirect obtained better performance in 5 target states (8-0, 9-1, 9-2, 11-0 and 14), comparable performance in 11 target states, and poorer performance in 2 target states (3 and 9-0). Taking state 8-0 as an example (Fig. 3b), scDirect identified NA5R2 and NA5R1 with the 5^th^ and the 6^th^ rank, respectively, while scDirect_WOE only identified NA5R2 with the 8^th^ rank. Besides, scDirect raised the ranks of key TFs like GRHL1, KLF4, EOMES and KLF17 (Supplementary Fig. 1). The improvement may be attributed to that the GAT model learned broader regulatory relations of key TFs.

Focusing on the scDirect identified TF list for each target state, each TF is given two biologically meaningful scores of the expected alteration and the directing score (Fig. 3b and Supplementary Fig. 1). The expected alteration represents the expression variation that the TF is supposed to change to direct the source state to the target state, and the directing score indicates how similar the perturbed cell state to target cell state after changing the TF with expected alteration, which is used to rank the TFs. The identified key TFs generally exhibited significantly higher directing scores than remaining TFs (Fig. 3b and Supplementary Fig. 1), which indicates their importance in directing cell state transitions. For instance, scDirect identified key TF LHX6 in target 16 with the highest expected alteration and directing score.

We further examined the target states in which scDirect exhibited unsatisfactory performance. In some target states both scDirect and baseline methods could not identify any ground-truth TF in the top 10 TF list, such as the state 11-1 and 13. We noticed that the ground-truth TF expression in these cases is either consistent between the source state and target state (Supplementary Fig. 2a) or especially low in the whole dataset (Supplementary Fig. 2b), and these gene expression patterns make it challenging to identify key TFs. In state 3 and 8-0, scDirect performed slightly worse than scDirect_WOE. We further compared the identification result of state 3 between scDirect and scDirect_WOE, and found that with the GAT enhancement, scDirect raised the rank of KLF1 from 42^nd^ to 34^th^ and the rank of KLF5 from 2^nd^ to 1^st^ while diminished the rank of KLF2 from 28^th^ to 35^th^ (Supplementary Fig. 3). Most cells of the state 3 expressed KLF1 and KLF5 with high values, while few cells expressed KLF2, which indicates that the GAT enhancement may ignore the key TFs with low expression in target states. Beyond the aforementioned cases, due to that some TFs lacks the binding motif in the databases, we missed it at the beginning of GRN inference. For example, in the state 9-0 the key TF GRHL3 is not included in the alternative TF list. To overcome such a limitation, we supplemented these TFs from existing knowledge databases of TFs regulating targets, such as NicheNet (Methods). We observed that with the modification scDirect could identified GRHL3 with the 24^th^ rank (Supplementary Fig. 4).

### 2.4 scDirect identifies TFs directing mouse hair follicle development

We then tried to demonstrate the reliability of scDirect in identifying key TFs in cell development. We collected a SHARE-seq dataset profiling mouse skin with paired scRNA-seq data and scATAC-seq data^23^. This dataset comprises cells of mouse hair follicle development system. In this system, Transit-Amplifying Cells (TACs) differentiate into different lineages: inner root sheath (IRS), hair shaft, or medulla. These lineages have been utilized to evaluate various computational methods^36,37^. Due to a large imbalance in cell types of original dataset, we subsampled equal number of each cell type in the hair follicle development system and obtained a dataset of 2,688 cells (Fig. 4a). 2,748 genes were retained after gene filtering. We applied scDirect on each lineage and selected top 10 TFs as the candidate TFs (Fig. 4b).

**Fig. 4.**
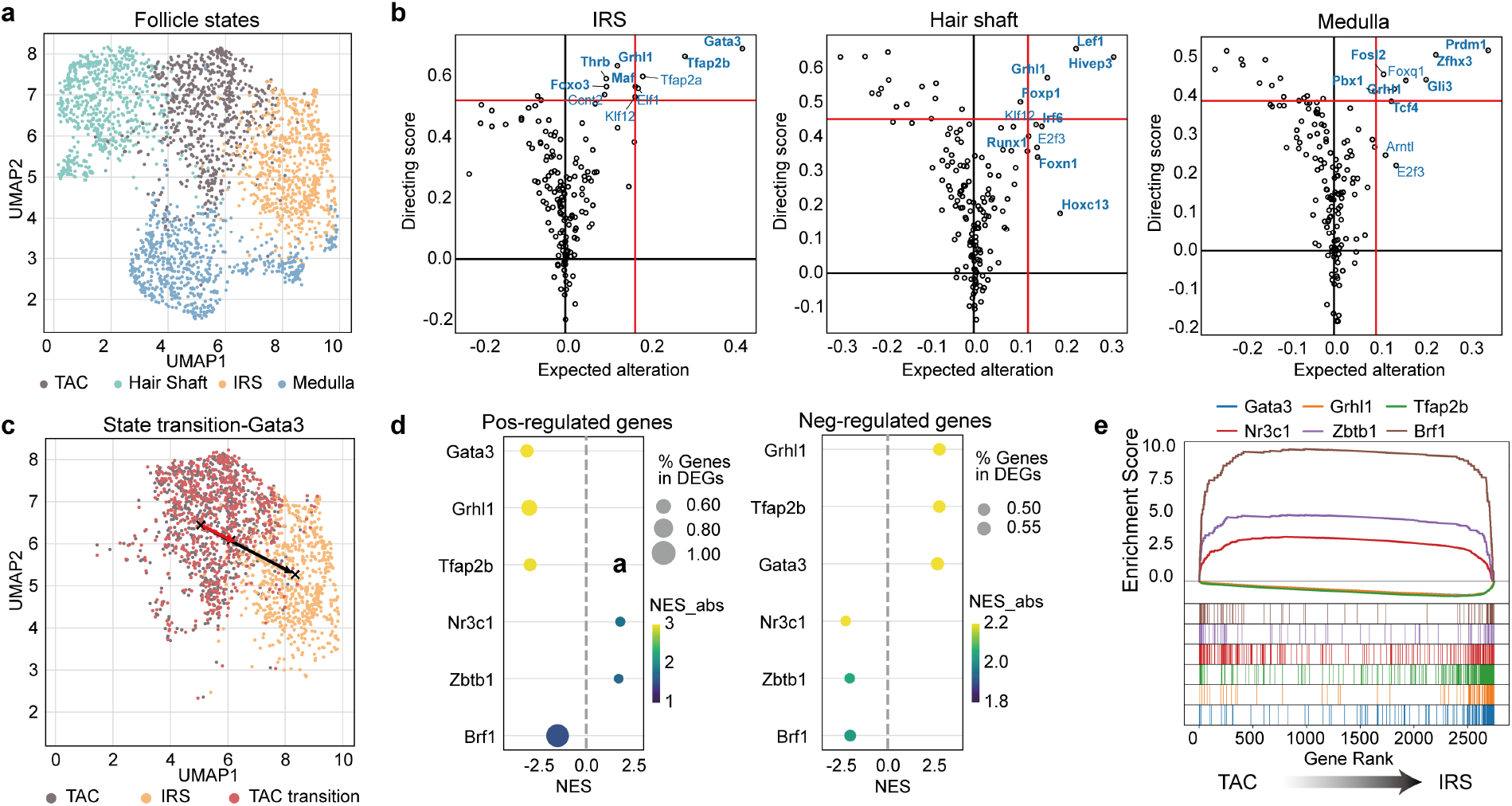
scDirect identifies key TFs in mouse hair follicle development. **a** UMAP visualization of down-sampled mouse hair follicle scRNA-seq data. **b** scDirect TF identification plot on mouse hair developing lineages of IRS, hair shaft, and medulla. Red lines are the quantile thresholds to filter 10 TFs. DEGs are annotated in bold. **c** UMAP visualization of TAC transiting towards IRS with Gata3 changing the expected variation. **d** Dot plots of gene set enrichment analysis (GSEA) results, showing enrichment of the positively and negatively regulated target sets of scDirect identified TFs within DEGs between TAC and IRS. TFs that rank top 3 and bottom 3 are selected. **e** Enrichment plot of TF positively regulated gene sets within DEGs between TAC and IRS.

In the IRS lineage, Gata3 is the top 1 identified TF with expected alteration 0.422 and directing score 0.691. It has been reported that Gata3 plays a direct role in differentially regulating cell lineages of hair follicle^38^, and loss of Gata3 negatively impacts the formation of IRS^39^. Our identification result is consistent with existing findings, and indicates Gata3 is a key TF directing the IRS lineage (Fig. 4c).

Among other identified TFs in the IRS lineage, Tfap2a is considered to be up-regulated in IRS during early hair follicle morphogenesis^40^, and Maf is also involved in hair morphogenesis of IRS layers^41^.

As for the hair shaft lineage, scDirect identified Lef1 as the top 1 TF. Lef1 is demonstrated as a key regulator of Wnt/β-catenin signaling in hair shaft differentiation ^42^. Besides, Runx1 is co-localized with Lef1 and is a specific marker of hair shaft^43^, and Hoxc13 is reviewed to affect hair shaft differentiation with various alternative mechanisms^44^.

Though medulla lineage has been less explored compared to other two lineages, we can still find some existing literature to support our identified TFs. For instance, the deletion of Prdm1 can result in aberrant medulla cell organization^45^, which indicates the important role of Prdm1 in medulla formation. Additionally, Foxq1 is discovered to control medulla differentiation through a common regulatory pathway as Hoxc13^46^. Even Foxq1 is not a differentially expressed gene (DEG), it can still be identified by scDirect, which suggests scDirect can identify key TFs despite the difference between target state and source state is relatively small.

Taking identified TFs of different lineages for comparative analysis, we found an interesting TF Grhl1 identified in all the lineages. Grhl1 is known to be dynamically expressed in the interfollicular epidermis (IFE) differentiation^47^, and is reported to associated with all the three lineages^48^, which is in agreement with our findings. We hypothesize Grhl1 may be an important regulator in hair follicle development. With considerable and reliable support of existing literature, we verified that scDriect can identify the key TFs in mouse hair follicle development.

We further characterized the identified TFs in detail taking the IRS lineage as a case. We investigated the relation between regulated targets of identified TFs and state DEGs. The 2,748 genes were ordered from TAC DEGs to IRS DEGs, and we divided regulated targets of each TF into positively regulated part and negatively regulated part according to the regulatory values. We conducted gene enrichment analysis (GSEA) to evaluate enrichment of the regulated target sets within TAC DEGs or IRS DEGs^49^. From the total ranked candidate TFs identified by scDirect, top 3 TFs (Gata3, Grhl1, and Tfap2b) and bottom 3 TFs (Nr3c1, Zbtb1, and Brf1) were selected. Target sets of these TFs were utilized to calculate normalized enrichment scores (NESs), which quantitatively measure the enrichment degree of gene sets. The NESs indicated the positively regulated target sets and negatively regulated target sets of top TFs are enriched in target IRS state and source TAC state, respectively (Fig. 4d), while regulated target sets of bottom TFs did not exhibit such enrichment profiles. Furthermore, we illustrated the enrichment result of positively regulated target set for each TF (Fig. 4e). For the top 3 TFs, most positively regulated targets were enriched in top IRS DEGs, while there was not a significant enrichment bias for bottom TFs regulated targets between the TAC and IRC state (Fig. 4e).

### 2.5 scDirect identifies the TFs directing cell reprogramming across different scenarios

The design of scDirect to identify TF combinations between cell state transitions could facilitate cell reprogramming experiments. We collected single-cell datasets for 5 different cell reprogramming scenarios of human and mouse: fibroblasts to keratinocytes^50^, fibroblasts to macrophages^51^, fibroblasts to cardiomyocytes^52^, and B cells to macrophages in mouse^53^, and fibroblasts to induced pluripotent stem cells (iPSCs)^2,54^,55 in human (Supplementary Fig. 5). Each reprogramming case has been extensively investigated with several reported key TFs (Fig. 5a). We denoted these cases from s1 to s5 for brevity (Fig. 5a). We compared scDirect and scDirect_WOE with D’Alessio method, Mogrify, and ANANSE, which are methods applied on bulk-omics data to identify key TFs. Mogrify prediction results are downloaded at https://mogrify.net/, and results of D’Alessio et al. and ANANSE are obtained from the original paper^13,17^. The methods that did not report the TF list of our reprogramming scenarios were excluded in the comparison.

**Fig. 5.**
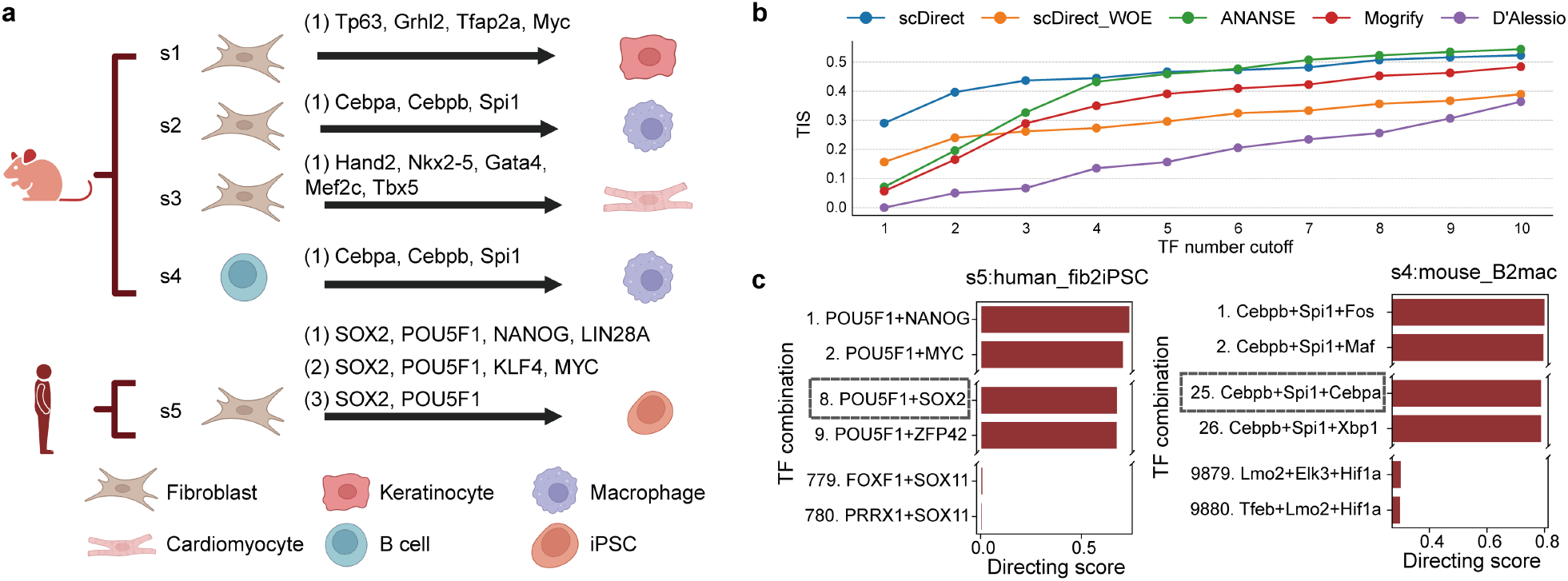
scDirect identifies the TFs directing cell reprogramming across different scenarios. **a** 5 reprogramming scenarios (s1-s5) with experimentally validated reprogramming TF combinations. **b** The line plots show the comparison of different methods based on the TIS calculated between identified TFs and ground-truth reprogramming TFs. **c** Ranked TF combinations with scDirect predicted directing scores. the ground-truth combinations of case s4 (Cebpa, Cebpb, and Spi1) and s5 (POU5F1 and SOX2) are marked with dashed boxes.

To comprehensively analyze the performance of methods, we calculated the average TIS across different scenarios for each method, and illustrated results with rank cutoff of identified TF list from 1 to 10. With TF rank cutoff from 1 to 3, scDirect outperformed other methods with significantly higher HISs. With TF rank cutoff from 4 to 10, scDirect and ANANSE achieve higher HIS than Mogrify and D’Alessio method, and the performance of scDirect and ANANSE is comparable. Moreover, scDirect significantly outperformed scDirect_WOE, which indicates that the GAT enhancement module effectively improves the identification result of reprogramming key TFs. Besides, scDirect identifies key TFs with the highest rank in 4 of 5 reprogramming cases (Supplementary Fig. 6). These results demonstrate that scDirect outperforms baselines and identifies key reprogramming TFs with high rank.

In addition to the better performance than existing methods, to the best of our knowledge, scDirect is the only method capable of quantitatively identifying possible TF combinations in cell state transition. For each combination of TFs, scDirect could calculate a directing score (Methods). We took reprogramming case s4 and s5 as examples. We applied scDirect to acquire a ranked list of 40 top TFs for each case, and then calculated and ranked all the possible combinations of 3 TFs and 2 TFs for case s4 and s5, respectively. As shown in Fig. 5c, the ground-truth combination of case s4 (Cebpa, Cebpb, and Spi1) ranks the 25^th^ in total 9,880 combinations, and the ground-truth combination of case s5 (POU5F1 and SOX2) ranks the 8^th^ in total 780 combinations. These results demonstrate that scDirect could assist and accelerate cell reprogramming experiments without any prior selection.

In wet lab experiments, candidate TFs for directing cell programming are sometimes hand-picked by prior knowledge or experimental feasibility. To further explore the potential of scDirect with hand-picked candidate TFs, we *in silico* repeated Yamanaka’s experiment^2^ which reprogramed fibroblasts to pluripotent stem cells with a combination of 4 TFs (POU5F1, SOX2, KLF4, and MYC). We added the 24 candidate TFs that Yamanaka et al. chose in their experiment, to our processed single cell dataset. There are 10,626 candidate combinations of selecting 4 TFs in the top 24 TF list, and scDirect identified Yamanaka factor combination with the 50^th^ rank (Supplementary Fig. 7), which suggests that if Yamanaka applied scDirect before conducting experiments, he would find the best combination in the 50^th^ experiment, which leads to a substantial decrease in both cost and time.

## 3 Discussion

In this paper, we presented scDirect, a computational framework to identify key TFs in state transitions. As far as we know, scDirect is the first method specially designed for identifying key TFs for directing cell state transitions at the single-cell level, and can be directly applied on single-cell datasets with either annotated cell types or unnamed clusters. scDirect quantitatively identifies key TFs between cell state transitions and assigns a directing score to arbitrary TF combination. We demonstrated that scDirect exhibit superior performance in cell differentiation, somatic cell conversion, and cell reprogramming.

Understanding cell state transitions could greatly contribute to biomedical research and drug discovery. Many researchers have been dedicated to discovering effective TF combinations directing cell state transitions^7,51,56^.The efforts on engineering cells towards expected states significantly facilitate regenerative medicine. scDirect can serve as an efficient tool to expedite these scenarios by providing candidate TFs as well as their combinations with high confidence.

We established a comprehensive benchmark framework for validating computational methods on TF identification. We utilized a TF-overexpression single-cell atlas [cite] and defined a benchmarking metric TIS to consider both rank and number of identified ground-truth TFs. Besides, we collected multiple single-cell datasets of experimentally validated reprogramming cases to confirm the applicability of scDirect in cell reprogramming. The benchmark framework and the single-cell datasets may benefit further studies on the development of related algorithms.

There are also several avenues for improving scDirect. Cell state transition is a continuous biological process, and thus the framework of scDirect can be extended to model TF variations as a changing tendency during the whole transition. Besides, not only key TFs but also some other regulation components, such chromatin regulators, splicing factors, and microRNAs^57^, can be incorporated into scDirect. In addition, though the linear model of scDirect has been proved to be efficient in this study, it is supposed to be valuable to incorporate with non-linear regulatory relations when dealing with more complex cell state transitions.

## 4 Methods

### 4.1 Data preprocessing

We process scRNA-seq data following the standard pipeline of Scanpy^27^. We select highly variable genes (HVGs) between source and target state cells with a fixed dispersion threshold. As the feature selection with HVGs would miss some important TFs, we further select TFs to supplement the gene set of interest. We calculate three sets of TFs: (1) Top 400 TFs highly variable in source and target cells; (2) Top 100 TFs differentially expressed between source and target cells; (3) Top 100 TFs highly expressed in target cells. We take the intersection of these sets as supplementary TFs and append them into the gene set of interest. The final gene set of interest usually includes 2,000 to 3,000 genes.

### 4.2 Primary GRN construction with CellOracle

As we model cell state transition as a linear process, we expect a GRN maintaining linear relations between TFs and target genes. Here we use CellOracle^25^ to construct linear GRNs, and it is proved to be more accurate than tree-based ensemble methods, such as SCENIC^58^ and GENIE3^59^. CellOracle takes both scRNA-seq data and scATAC-seq data as input.

Following CellOracle workflow, we first use the whole cells of scATAC data to construct a base GRN matrix 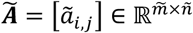, where 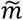 and *ñ* denote number of targets and number of TFs, respectively. Note that target here means a gene that is regulated by a specific TF, and may also be a TF itself. *ã*_*i,j*_ ∈ [0,1] represents whether TF *j* has a regulatory relation with gene *i*, and 1 means yes. CellOracle locates transcriptional start sites (TSSs) within the accessible chromatin regions, which are also called peaks, then applies Cicero^60^ to identify the correlated peaks to the TSS for each gene. Then the DNA sequence of each correlated peak is used to scan TFs with the gimmemotifs v.5 vertebrate motif dataset^61^. For the TFs that not included in the gimmemotifs database, we search the corresponding motifs in JASPAR database^62^. In this way the TFs and targets can be linked, and after regulatory score filtering we acquire the base GRN matrix ***Ã*** The calculation of ***Ã*** follows the default parameters of CellOracle. If a TF misses the motif sequence in the motif databases and it is required as a candidate TF, we link it with the targets based on the existing knowledge database NicheNet to supplement ***Ã***.

Then Bagging Ridge model is applied to predict the expression of a target gene based on regulatory TFs identified in *Ã* and the output is a posterior distribution of coefficient value *â*_*i,j*_:

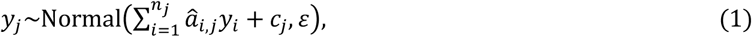

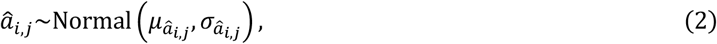

where *y*_*j*_ is the single target gene expression, *y*_*j*_ is candidate TF expression, *n*_*j*_ is the number of candidate TFs regulating *j*th gene, and *c*_*j*_ is the intercept. 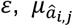 and 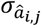 are parameters of normal distributions. Then we get the primary GRN matrix ***Â*** = [*â*_*i,j*_]∈ ℝ^*m*×*n*^, where *m* and *n* denote number of total genes and number of TFs, respectively, and *â*_*i,j*_ represents the regulatory coefficient of *j*th TF to *i*th target. Note that ***Â*** is acquired from scRNA-seq data based on the positive links in ***Ã***, so the size of ***Â*** is generally less than ***Ã***. For each transition from a source cell state to a target cell state we calculate a primary GRN matrix ***Â***, and the calculation of ***Â***follows the default parameters of CellOracle.

### 4.3 GRN enhancement with graph attention networks (GAT)

As shown in Fig. 1b, the GAT prediction model consists of two parts, the GAT encoder and the multilayer perceptron (MLP) predictor. The input of the encoder is the node feature matrix ***X*** ∈ ℝ^*c,m*^ and the adjacent matrix ***M*** ∈ ℝ^*m*×*m*^, where *c* is the total number of source and target cells. ***X*** is the processed log-normalized scRNA-seq expression matrix, and ***M*** is transformed from ***Â*** by connecting each TF-target pair whose value is not zero in ***Â*. *X*** and ***M*** comprise the input graph. In the input graph, each node is a gene represented by a vector ***x***_,***j***_ ∈ ℝ^*c*^, a column of ***X***. The encoder consists of three graph attentional layers, each layer learns a shared weight matrix 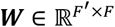 and takes a set of node features as input, ***G*** = {***g***_1_, ***g***_2_, …, ***g***_*m*_}, ***g***_*i*_ ∈ ℝ^*F*^, and produces a new set of node features, 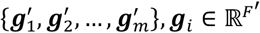 where *F* and *F*^**′**^ are feature numbers. In the first layer, *F* is equal to *c*. The attention coefficient of gene *j* to gene *i* is calculated by:

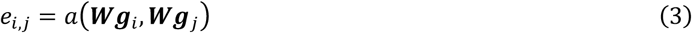

where the attention mechanism *a* is a single layer feedforward network. Only neighbors of gene *i* are used to calculate the attention coefficients, and then the coefficient is normalized by the Softmax function:

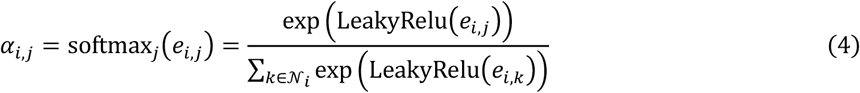

where **𝒩**_*i*_ is the first-order neighbors of gene *i*, including *i*, and LeakyRelu is a non-linear activation function. Multi-head attention mechanism is then applied to stabilize the learning process, and the output of gene *i* is:

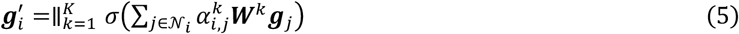

where ∥ is concatenation operation, and *σ* is the ELU activation function. For the last graph attention layer, the concatenation is replaced by averaging:

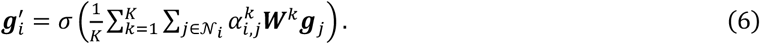

After the GAT encoder, the gene *i* and gene *j* are encoded to ***ĝ***_*i*_ and ***ĝ***_*j*_, respectively. The predicted score of gene *i* regulating gene *j* is calculated by:

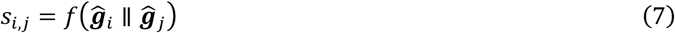

where *f* is a three-layer MLP. Here we denote ***L*** = [*l*_*i,j*_] ∈ ℝ^*m*×*n*^ as a binary label matrix indicating the regulatory relations between TFs and targets, and L is transformed from ***Â*** by converting all the non-zero values in ***Â*** to 1. Here we use the binary cross entropy loss to optimize the model:

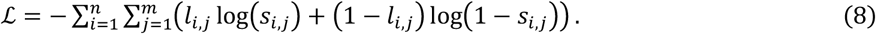

The details of model training and predicted link filtering are shown in Supplementary Note 1. After adding the recovered links, we acquire an enhanced GRN matrix ***A*** ∈ ℝ^*m*×*n*^.

### 4.4 Model training of graph attention networks (GAT)

The label matrix ***L*** is generally class-imbalanced and the negative links are significantly more than positive links. To handle the problem, we follow the data split strategy of DeepTFni^31^ using a tenfold scheme. We split the positive links into ten subsets of equal size. One subset of which and a same number set of randomly sampled negative links form the test set, and all the remaining links form the train set. The splitting repeats ten times, and each set of the positive links is used for one time as a test set.

The model is built by the Pytorch library and DGL library. The initial learning rate is 0.01, and Adam optimizer is used to optimize the model. The maximum epoch number is 1500, and early stopping strategy is adopted. The head numbers of the GAT layers are fixed to 4, 4, and 6, respectively. The hidden GAT layer dimension is set to 16, and the output GAT layer dimension is set to 7.

After the model is trained, for each fold we set the median predicted value of the test set as the classification threshold. We use the trained model to predict each TF-target link and assign a value, then these values are classified to 0 and 1 based on the threshold, and 1 represents the link is predicted as positive. The links predicted as positive in more than 9 folds are retained as candidate recovering links. Among these candidate links, top 5% links are further selected to be the final recovering links based on the predicted scores to discard the false positive samples.

Then we set the final recovering links as positive in ***Ã*** and repeat the CellOracle^25^ GRN calculation process of getting ***Â***, to acquire an enhanced GRN matrix ***A*** ∈ ℝ^*m*×*n*^.

### 4.5 Modelling TF identification as a linear inverse problem

Suppose the transition is from source cell state *𝒮* to target cell state *𝒯*. The processed gene expression data of *𝒮* is 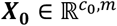 and processed gene expression data of *𝒯* is 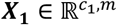 where *c*_0_ and *c*_1_ are the cell number of *𝒮* and *𝒯*, respectively. We then average the expression across all the cells of *𝒮* and *𝒯*, respectively, to acquire mean expression vectors ***y***_0_ ∈ ℝ^*m*×1^ and ***y***_**1**_ ∈ ℝ^*m*×1^. The state transition vector can be calculated by:

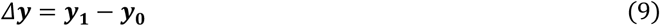

where *Δ****y*** indicates the direction that governs the cell state transition. We assume the alteration of TFs will direct cells to deviate from the original state by influencing the regulated target genes. With a linear GRN matrix *A*, we can model the transition process by the regulating relations of TFs to targets:

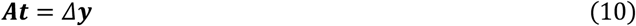

where ***t*** ∈ ℝ^*n×*1^ represents the variations of TFs. The solving of ***t*** with known ***A*** and *Δ****y*** is commonly referred to as a linear inverse problem ^63^. This equation describes the state transition that TFs only transmit one-step signals to the regulated targets, while in the GRN the TFs can propagate regulatory signals with multiple iterations. Following the setting of CellOracle, we set the propagation iterations to 3, and the GRN matrix is modified as:

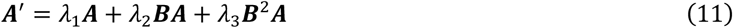

where ***B*** = [*b*_*i,j*_] ∈ ℝ^*m*×*n*^ is the GRN matrix transformed from ***A***, for gene *j* that regulates gene *i* the *b*_*i,j*_ is set as the value in ***A***, and otherwise is set to 0. *λ*_1_, *λ*_2_ and *λ*_3_ denotes the weights of TF propagation of one-step, two-step and three-step, respectively. Considering the one-step propagation of TFs predominantly influence the regulatory process, in this paper we empirically fixed *λ*_1_, *λ*_2_ and *λ*_3_ to 0.6, 0.2 and 0.2, respectively. In our case, the row number *m* of ***A***^**′**^ is larger than the column number *n*, which leads to an over-determined problem and may not have a solution. Consequently, we aim to find a 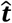 that ensures 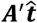 closely resembles *Δ****y***. Additionally, we add the Tikhonov regularization to restrict the range of 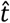, and it can be solved by:

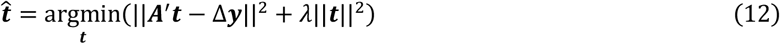

where *λ*||***t***||^2^ is a Tikhonov regularization, also called L2 regularization, and *λ* controls the degree of the regularization. We apply the ridge regression function in the scikit-learn library ^64^ to solve the problem, and *λ* is set to 1. The solved 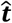 represents the expected alterations of TFs.

After acquiring the expected alteration vector 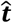, for each TF we calculate a directing score to measure the importance in the transition procedure. For each TF, keep the corresponding value and set other values to 0 in 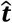 to get 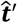 and the Pearson correlation coefficient (PCC) between 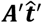 and Δ***y*** is calculated as the directing score. The final TF set is selected based on both expected alterations and directing scores with a same quantile threshold, and ranked by directing scores. In order to acquire a candidate TF list with a specific number, the quantile threshold is calculated with iterative searching. The directing score of a TF combination could be calculated in a same way.

### 4.6 Benchmark construction on a TF-overexpression hESC atlas

Joung et al. created a barcoded open reading frame (ORF) library of 3,548 TF splice isoforms^29^. The barcoded TF ORFs were packaged into lentivirus and then transduced into hESCs at low multiplicity of infection (MOI) to perform single TF overexpression across cells. These hESCs were then profiled by scRNA-seq to get a TF overexpression single-cell atlas. After down sampling, the atlas retains 671,453 cells covering 3,266 TFs. The original atlas is clustered to 9 major clusters. Clusters 6-8 are confirmed to be differentiated cells and were further subclustered to 25 minor clusters^29^. The major cluster 5 is the most adjacent cluster to the differentiated cells in the original low-dimensional space, and we assume it as the source cell state to be converted to those differentiated cells. Then we randomly sampled 2,000 cells in major cluster 5 and combined them to the 25 minor clusters to form the benchmarking dataset.

The 2,000 sampled cells are regarded as the source cell state that possibly transits to the 25 target cell states (Supplementary Fig. 8). Each target cell state comprises distinct groups of overexpressed TFs, and these TFs are considered as ground-truth key TFs directing differentiation from the source cell state to the target cell states. For each TF in each target cell state, we divide the number of cells that indicate the TF ORF in the state by number of cells that indicate this TF ORF in the whole atlas, to calculate the percentage. Then we filter out the TFs of percentage less than 5% and counting cells less than 5 in each target cell state, and 15 target cell states are retained with key TFs counting from 1 to 8 (Fig. 3a).

Then for each target cell state we define a TIS between the ground-truth key TF list **𝒦** and identified candidate ranked TF list **𝒞**:

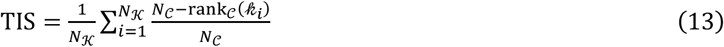

where *N*_**𝒦**_ is length of **𝒦**, *N*_*𝒞*_ is the length of *𝒞*, and *𝓀*_*i*_ is the *i*th TF in *𝒦*. If *𝓀*_*i*_ is in *𝒞, rank*_*𝒞*_(*𝓀*_*i*_) means the rank of *𝓀*_*i*_ in *𝒞*, and the rank starts from zero not including the former elements in *𝒞*. For example, if top three elements in *𝒞* is *𝓀*_1_, *c*_1_, *c*_2_, and *𝓀*_1_ (*c*_1_ and *c*_2_ are not in **𝒦**), and *rank*_*𝒞*_(*𝓀*_1_) and *rank*_*𝒞*_(*𝓀*_2_) are 0 and 2, respectively. For the *𝓀*_*i*_ not in *𝒞*, the *rank*_*𝒞*_(*𝓀*_*i*_) will be *N*_*𝒞*_. Consequently, TIS can be a comprehensive metric ranging from 0 to 1 and considering both the ranks and numbers of identified TFs.

## Supporting information

All the supplementary files

## Data Availability

In this paper we used a series of single-cell datasets, mainly including the scRNA-seq and scATAC-seq datasets. Every scRNA-seq dataset is accompanied with a scATAC-seq dataset, and ensure both are from the same samples or the homologous tissues. Details and accessions of the datasets are listed in Supplementary Table 1. The computational tools used for data processing and analysis are summarized in Supplementary Table 2.

## Code Availability

The code of scDirect is available at https://github.com/Chen-Li-17/scDirect.

## Acknowledgements

The work is supported in part by National Natural Science Foundation of China (grants 62373210, 62250005, 61721003), and National Key R&D Program of China (grant 2021YFF1200900).

## Author contributions

C.L. and L.W. conceived the main idea of the study. C.L. designed and developed the method. C.L. performed the data analysis and method benchmarking. S.C., Y.C. and M.H. contributed to the discussion of results. H.B. helped with the construction of the method framework. L.W. supervised the study. C.L., L.W. and X.Z. wrote the manuscript with the help from other authors.

