## Supplementary material for "scDirect: key transcription factor identification for directing cell state transitions based on single-cell multi-omics data": All the supplementary files

### Supplementary Information for

### scDirect: key transcription factor identification for directing cell state transitions based on enhanced single-cell multi-omics gene regulatory networks

Chen Li[1], Sijie Chen[1], Yixin Chen[1], Haiyang Bian[1], Minsheng Hao[1], Lei Wei[1,*] and Xuegong Zhang[1,2]

[1] Ministry of Education Key Laboratory of Bioinformatics, Bioinformatics Division at the Beijing National Research Center for Information Science and Technology, Center for Synthetic and Systems Biology, Department of Automation, Tsinghua University, Beijing 100084, China

[2] Center for Synthetic and Systems Biology, School of Life Sciences and School of Medicine, Tsinghua University, Beijing 100084, China

### Supplementary Figures

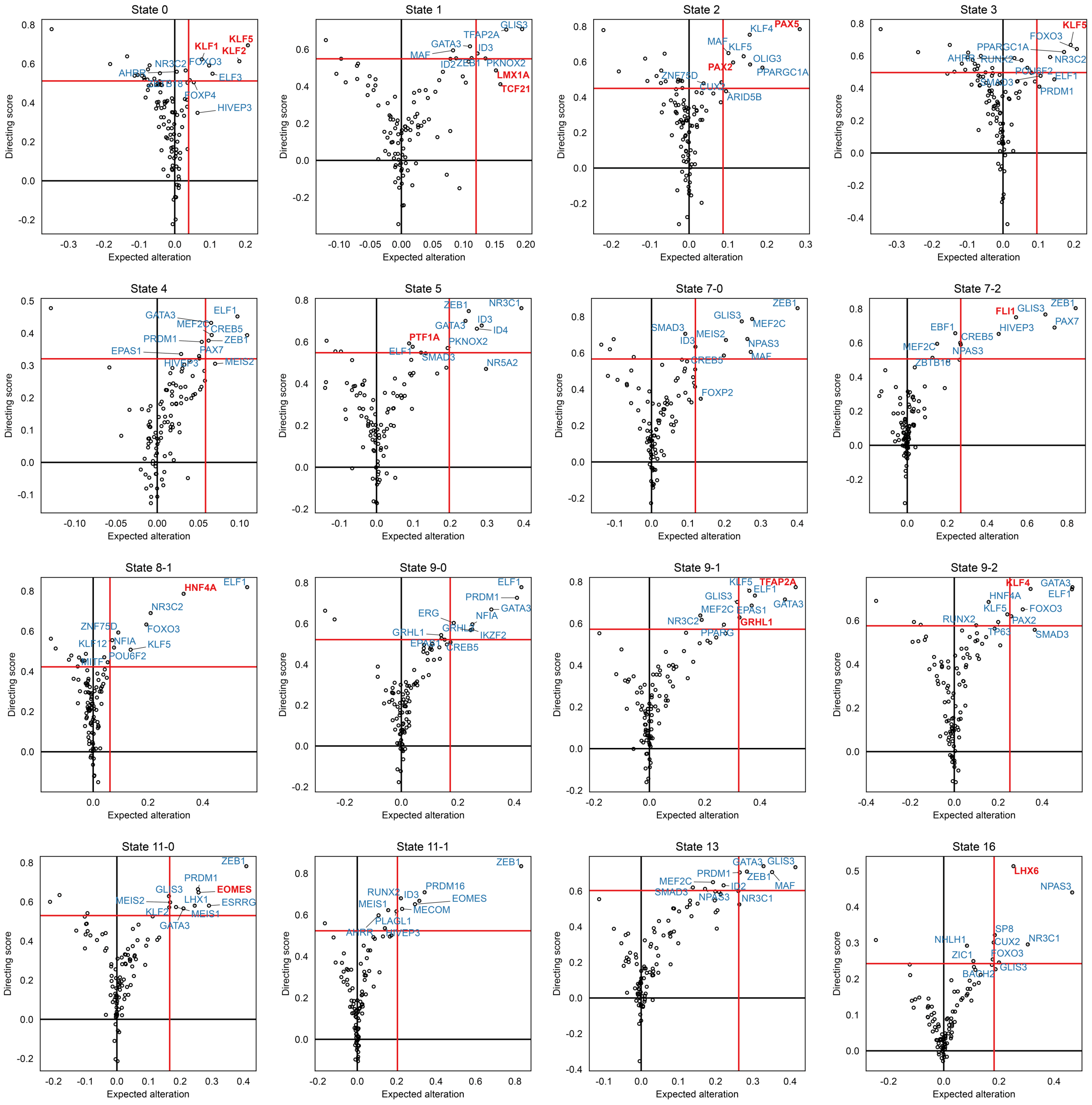

**Supplementary Fig. 1.** scDirect TF identification plot on other target cell states. Red lines are the quantile thresholds to filter 10 TFs. Key TFs are annotated in red.

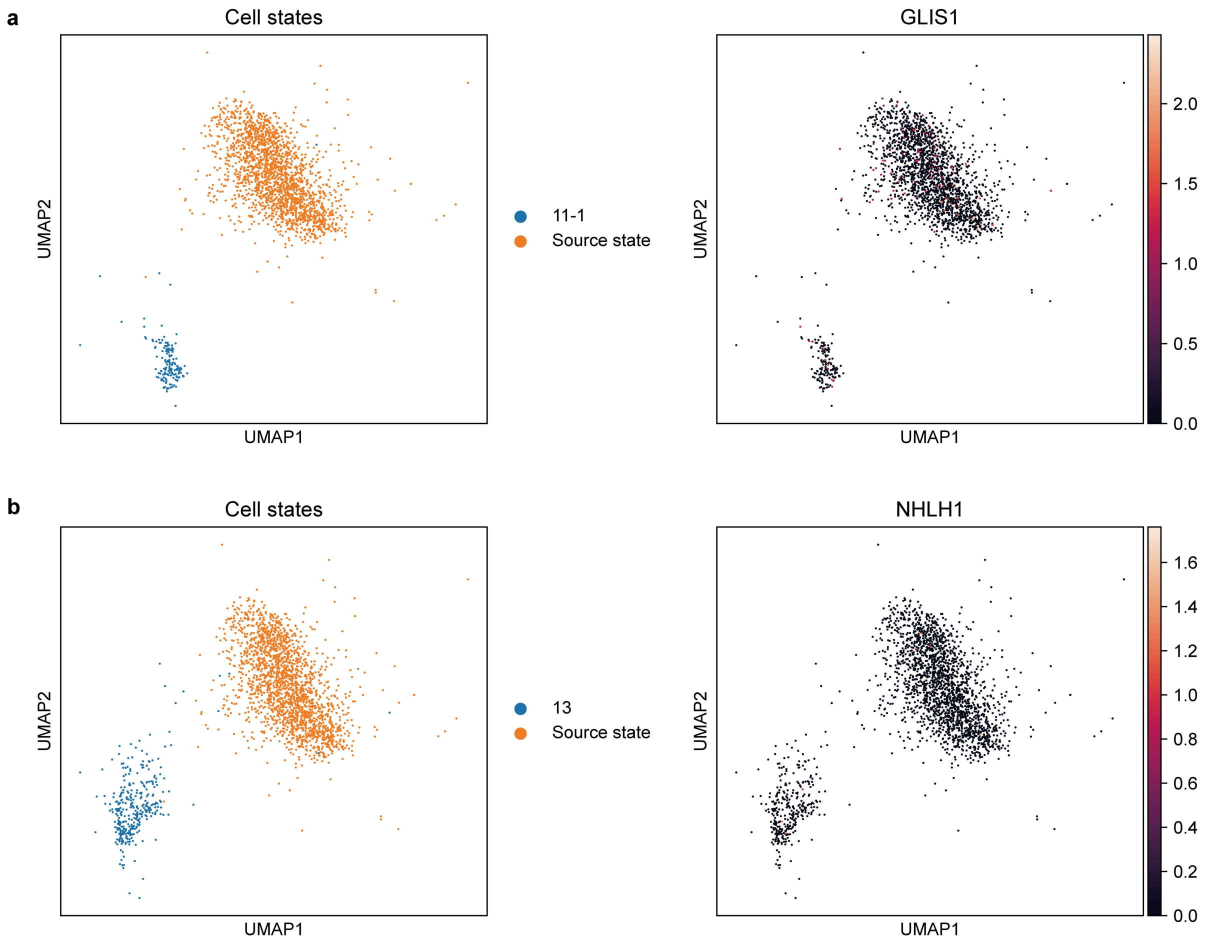

**Supplementary Fig. 2. a** The UMAP visualization of the source state and the target state 11-1. The normalized expression of key TF GLIS1 is shown on the UMAP embeddings. **b** The UMAP visualization of the source state and the target state 13. The normalized expression of key TF NHLH1 is shown on the UMAP embeddings.

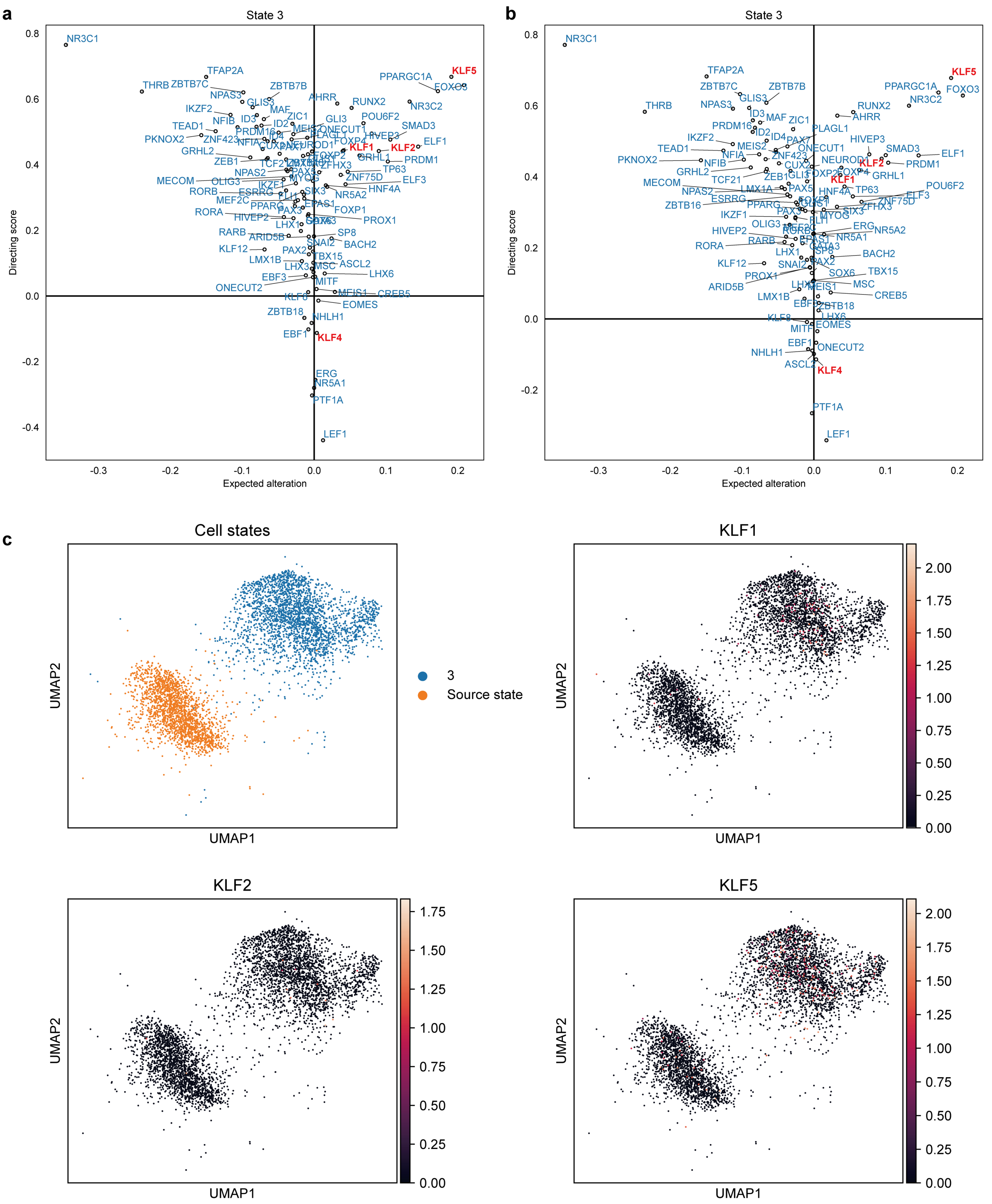

**Supplementary Fig. 3. a** scDirect TF identification plot on target state 3. Key TFs are annotated in red. **b** scDirect_WOE TF identification plot on target state 3. Key TFs are annotated in red. **c** The UMAP visualization of the source state and the target state 3. The normalized expression of key TF KLF1, KLF2 and KLF5 is shown on the UMAP embeddings.

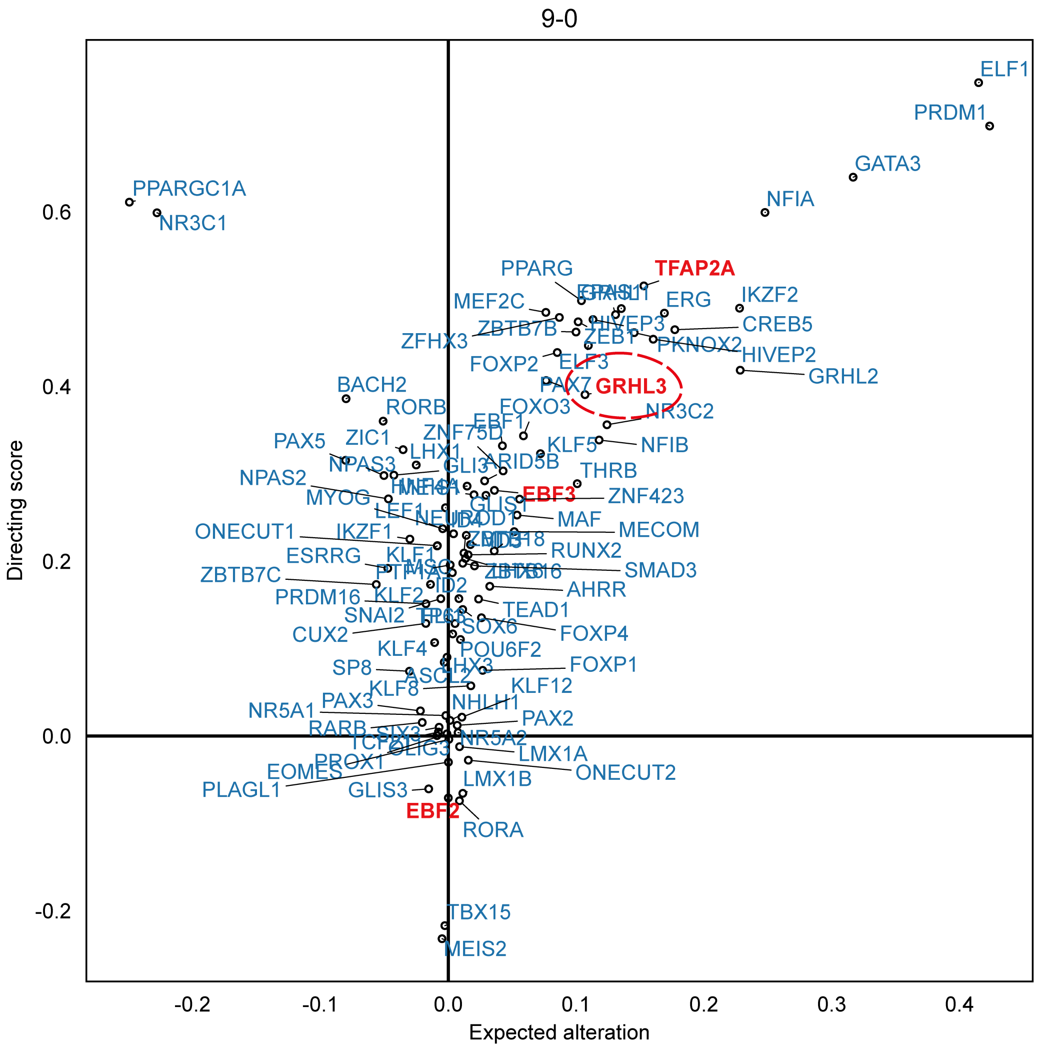

**Supplementary Fig. 4.** scDirect TF identification plot on target state 9-0. The knowledge of NicheNet database is added to supplement the TF-target links. Key TFs are annotated in red. The added key TF GRHL is annotated in a red circle.

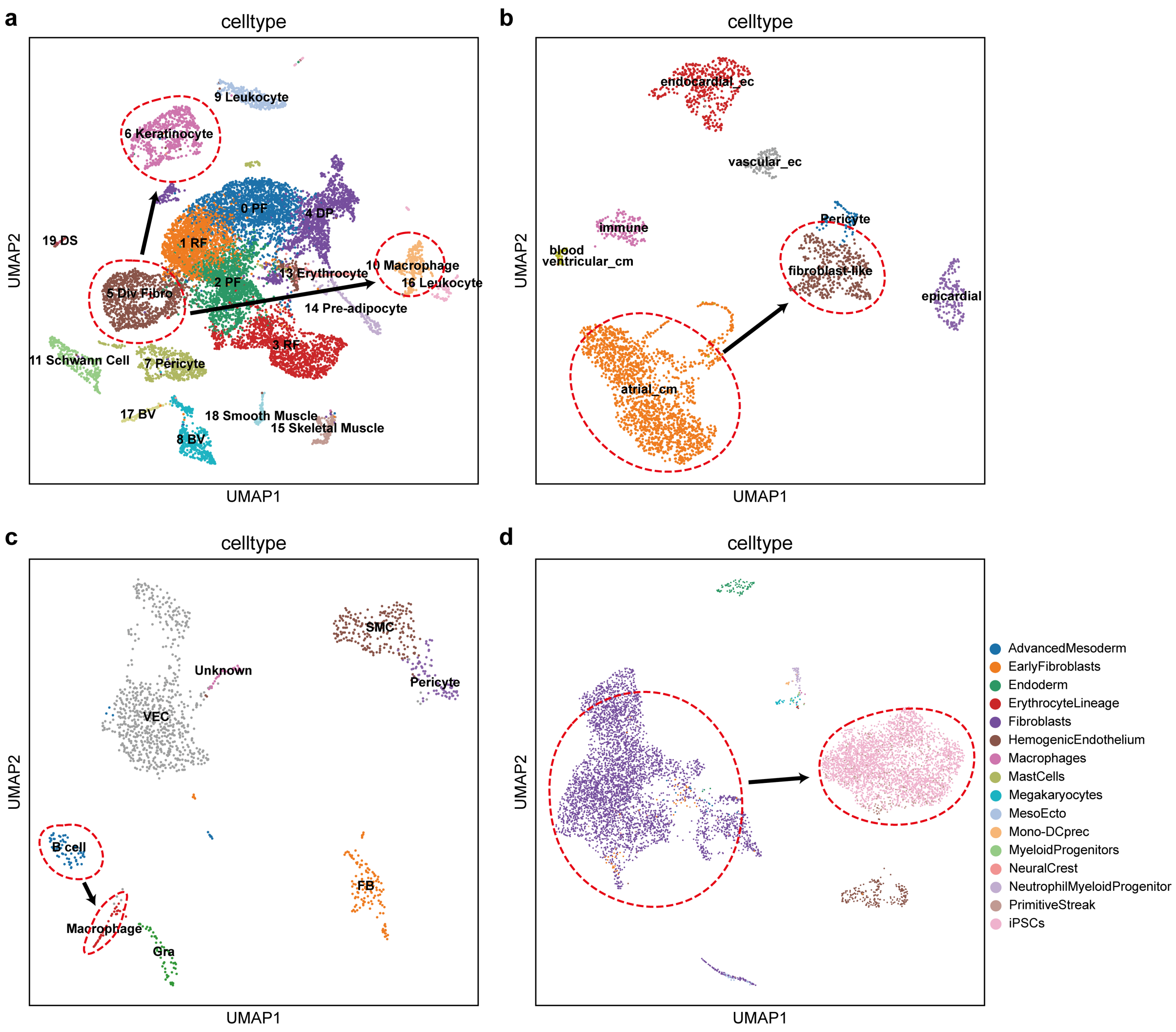

**Supplementary Fig. 5.** UMAP visualization of single-cell datasets for reprogramming cases. The dashed red circles represent source or target cell states. The black arrows point from source cell states to target cell states. (a) Fibroblasts to keratinocytes or macrophages in mouse. (b) Fibroblasts to cardiomyocytes in mouse. (c) B cells to macrophages in mouse. (d) Fibroblasts to induced pluripotent stem cells (iPSCs) in human.

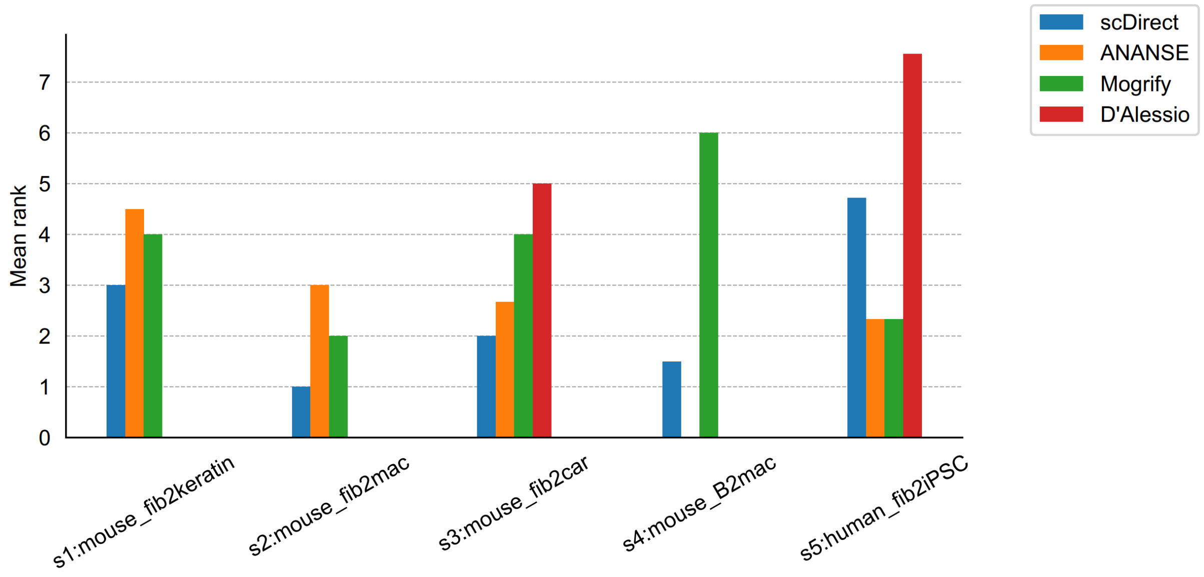

**Supplementary Fig. 6.** TF identification comparison by mean rank across different reprogramming cases.

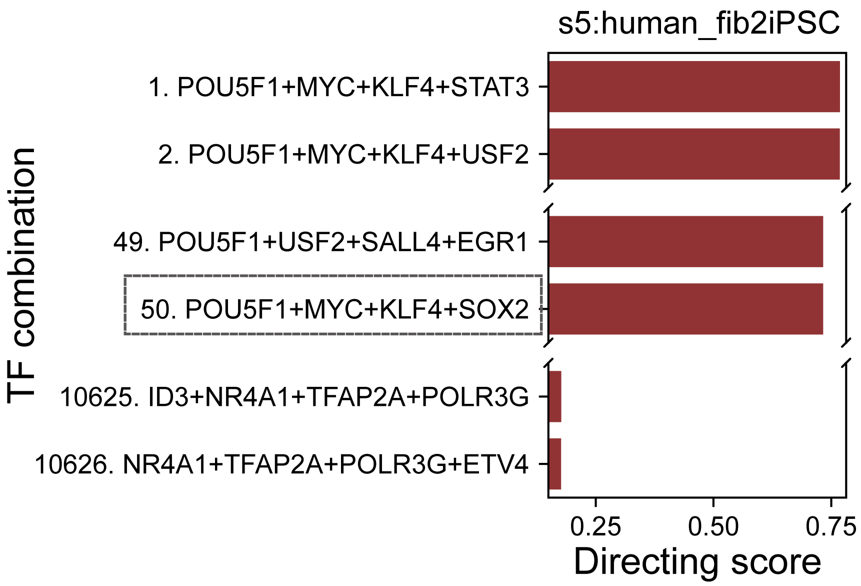

**Supplementary Fig. 7.** xxxxxx

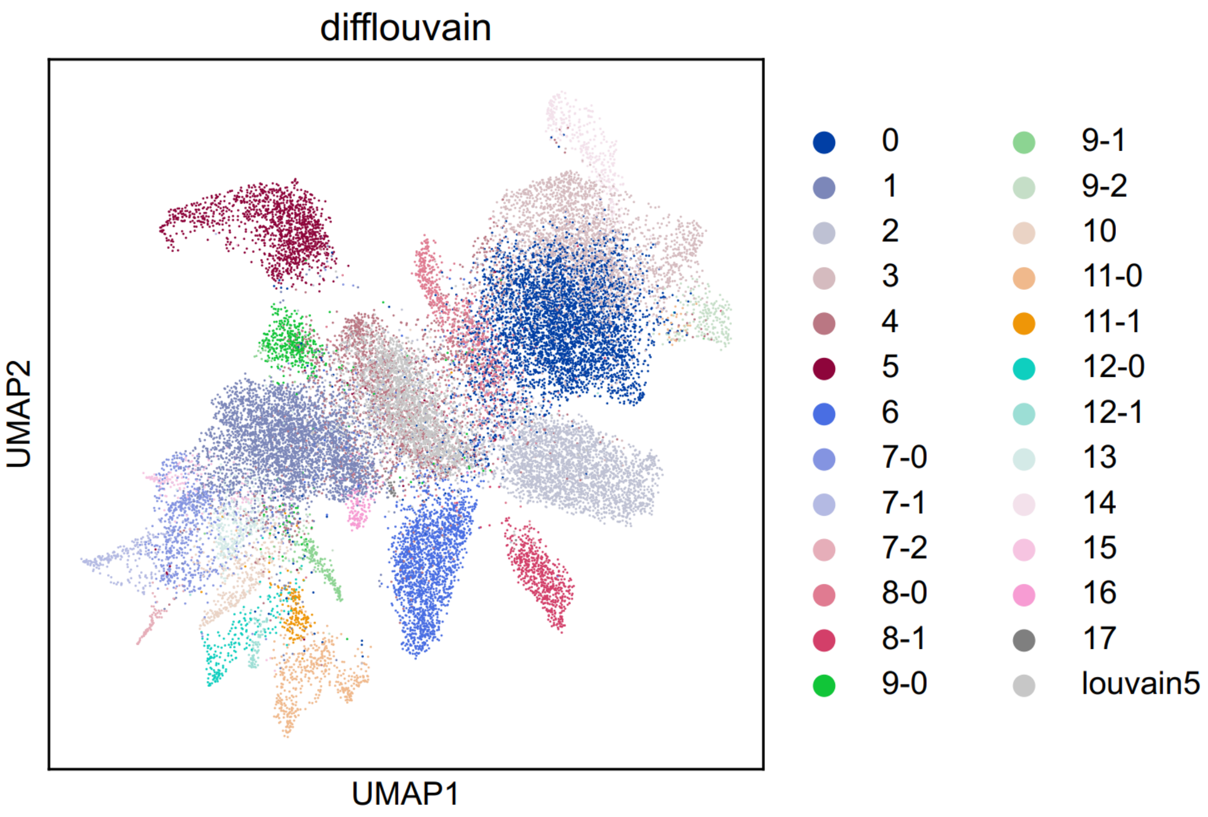

**Supplementary Fig. 8.** UMAP visualization of 2,000 sampled cells and 25 original target cells.

The ‘louvain5’ is the cluster of sampled cells.

#### Supplementary Tables

**Supplementary Table 1.** Summary of datasets used in this study.

| | scRNA | | | scATAC | | |
|---|---|---|---|---|---|---|
| Description | Tissue | Data accession | Reference | Tissue | Data accession | Reference |
| Data for TF identification benchmark. | Human differentiated part of hESCs | GSE217215 | [1] | Human brain | GSE174367 | [2] |
| Data for identifying TFs in mouse hair follicle development. | Mouse skin | GSE140203 | [3] | Mouse skin | GSE140203 | [3] |
| Data for reprogramming case of fibroblasts to keratinocytes in mouse. | Mouse neonatal skin | GSM5696148 | [4] | Mouse neonatal skin | GSM5696149 | [4] |
| Data for reprogramming case of fibroblasts to macrophages in mouse. | Mouse neonatal skin | GSM5696148 | [4] | Mouse neonatal skin | GSM5696149 | [4] |
| Data for reprogramming case of fibroblasts to cardiomyocytes in mouse. | Mouse neonatal heart | GSM5795776 | [5] | Mouse neonatal heart | GSM4644946 | [6] |
| Data for reprogramming case of B cells to macrophages in mouse. | Mouse neonatal heart | GSM4644956 | [6] | Mouse neonatal heart | GSM4644948 | [6] |
| Data for reprogramming case of fibroblasts to iPSCs in human. | iPSCs differentiation into macrophages | https://www.hipimmuneatlas.org/ | [7] | iPSCs differentiation into macrophages | E-MTAB-11616 | [7] |

**Supplementary Table 2.** Summary of computational tools.

| Method | Description | Version |
|---|---|---|
| Scanpy | Used for processing single-cell datasets. | 1.9.3 |
| Seurat | Used for processing single-cell datasets. | 4.1.1 |
| ArchR | Used for processing scATAC datasets. | 1.0.1 |
| CellTypist | Used for cell annotation. | 1.6.0 |
| MACS2 | Used for peak calling. | 2.2.7.1 |
| CellRangerATAC | Used for transforming fastq files to fragment files. | 2.1.0 |
| PyTorch | Used for training deep learning model. | 1.12.1 |
| DGL | Used for constructing graph neural networks. | 1.1.1 |
| scikit-learn | Used for performing ridge regression. | 1.2.2 |
